# Sex Differences in Aggression: Differential Roles of 5-HT_2_, Neuropeptide F and Tachykinin

**DOI:** 10.1101/407478

**Authors:** Andrew N. Bubak, Michael J. Watt, Kenneth J. Renner, Abigail A. Luman, Jamie D. Costabile, Erin J. Sanders, Jaime L. Grace, John G. Swallow

**Affiliations:** Department of Neurology, University of Colorado-Anschutz Medical Campus; Center for Brain and Behavior Research, Basic Biomedical Sciences, University of South Dakota; Biology Department, University of South Dakota; Department of Integrative Biology, University of Colorado-Denver; Department of Biology, Bradley University

**Author notes:** Corresponding Author: John G Swallow.

**Keywords:** 5-HT2, Serotonin, Neuropeptide F, Tachykinin, RNAi, Aggression, Invertebrate, Social Isolation

## Abstract

Despite the conserved function of aggression across taxa in obtaining critical resources such as food and mates, serotonin’s (5-HT) modulatory role on aggressive behavior appears to be largely inhibitory for vertebrates but stimulatory for invertebrates. However, critical gaps exist in our knowledge of invertebrates that need to be addressed before definitively stating opposing roles for 5-HT and aggression. Specifically, the role of 5-HT receptor subtypes are largely unknown, as is the potential interactive role of 5-HT with other neurochemical systems known to play a critical role in aggression. Similarly, the influence of these systems in driving sex differences in aggressive behavior of invertebrates is not well understood. Here, we investigated these questions by employing complementary approaches in a novel invertebrate model of aggression, the stalk-eyed fly. A combination of altered social conditions, pharmacological manipulation and 5-HT_2_ receptor knockdown by siRNA revealed an inhibitory role of this receptor subtype on aggression. Additionally, we provide evidence for 5-HT_2_’s involvement in regulating neuropeptide F activity, a suspected inhibitor of aggression. However, this function appears to be stage-specific, altering only the initiation stage of aggressive conflicts. Alternatively, pharmacologically increasing systemic concentrations of 5-HT significantly elevated the expression of the neuropeptide tachykinin, which did not affect contest initiation but instead promoted escalation via production of high intensity aggressive behaviors. Notably, these effects were limited solely to males, with female aggression and neuropeptide expression remaining unaltered by any manipulation that affected 5-HT. Together, these results demonstrate a more nuanced role for 5-HT in modulating aggression in invertebrates, revealing an important interactive role with neuropeptides that is more reminiscent of vertebrates. The sex-differences described here also provide valuable insight into the evolutionary contexts of this complex behavior.

**Significance Statement:** Serotonin’s (5-HT) modulatory role in aggression is generally reported as inhibitory in vertebrates but stimulatory in invertebrates. Using a novel invertebrate model system, we provide evidence of common pathways of aggression at the 5-HT receptor subtype level as well as 5-HT’s interactive role with other neurochemical systems namely neuropeptide F and tachykinin. Additionally, we found that these effects were sex-dependent as well as stage-dependent affecting either the initiation or escalation stage of an aggressive contest. Our results reveal the impressive level of conservation with respect to neurochemical mechanisms among species as diverse as vertebrates and invertebrates, and highlights the need to consider multiple factors when determining potential taxonomic differences in how 5-HT mediates aggression.

## 1. Introduction

Serotonin (5-HT) appears to promote aggression in invertebrates (1,2), in contrast to the largely inhibitory effect seen in vertebrates (3, but see 4). Much of the empirical support for this dichotomy comes from studies using arthropod invertebrates, with increased expression of overt aggressive behavior and greater willingness to engage in conflict seen in decapod crustaceans (5–8), crickets (9), ants (10,11), and dipteran flies (12–15) following pharmacological or genetic elevations of 5-HT at the systemic level. While these findings support the presumption that 5-HT has opposing effects on invertebrate aggression from vertebrates, there are critical gaps in knowledge that need to be considered before accurately stating that 5-HT exclusively modulates invertebrate aggression in a positive manner.

A more nuanced role for 5-HT in invertebrate aggression emerges when considering involvement of receptor subtypes. In vertebrates, differential binding of specific 5-HT receptors, predominantly 5-HT_1A_, 5-HT_1B_, and 5-HT_2_ subtypes, has profound implications for aggressive behavior (16.17). Notable sequence and functional homology for these subtypes have been described in invertebrates (18), and the limited information available suggests some similarity in their influence on insect aggression (19). For example, specific pharmacological activation of 5-HT_2_ receptors has an anti-aggressive effect in both rodents (20) and *Drosophila* (29), suggesting 5-HT_2_ receptor function is conserved evolutionarily. In contrast, a divergent role is indicated for 5-HT_1A_ receptors, activation of which largely dampens mammalian aggression (21) while enhancing aggressive behavior in *Drosophila* (19). Whether these same similarities and differences in subtype function exist in invertebrates other than *Drosophila* remains to be determined.

It is also possible that 5-HT receptors have distinct functions in mediating the contextual expression of specific aggressive behaviors and their intensity, which in turn will direct how the conflict proceeds (i.e., initiation, escalation, and termination). For instance, while 5-HT_1A_ and 5-HT_2_ receptors are generally inhibitory for vertebrate aggression, agonists of these receptors can promote high intensity aggression in mammals during certain situations such as maternal, territorial, and self-defense (21,22), demonstrating these subtypes can exert opposing effects according to context. In male *Drosophila*, overall aggression is heightened or decreased according to 5-HT_1A_ or 5-HT_2_ receptor activation, respectively, but 5-HT_1A_ predominantly affects expression of low intensity aggression (threat displays) while 5-HT_2_ mediates high intensity behaviors such as lunging (19). Contextual modulation of aggression by 5-HT is also seen in male stalk-eyed flies (*Teleopsis dalmanni*). Specifically, smaller males display low intensity aggression and initiate less contests when faced with a larger opponent, but pharmacologically increasing 5-HT causes the smaller male to initiate and escalate fights by promoting high-intensity aggression (23). Interestingly, contest duration remains the same whether the smaller male is treated with 5-HT or not (23) In contrast, when contestants are of equal size, both aggression intensity and contest duration appear to be modulated as a function of the difference in brain 5-HT between opponents (10). In other words, size-matched opponents with closer brain 5-HT concentrations will engage in prolonged high intensity contests regardless of whether 5-HT has been increased in one male, with exogenous manipulation only making contests shorter and less intense if it causes brain 5-HT to be substantially elevated above that of the opponent (10). Thus, aggression in *T. dalmanni* provides a useful model for examining how 5-HT can discretely modulate behavioral expression according to specific contexts, which will in turn determine when conflicts are initiated/terminated and if there is an escalation in the intensity of aggression during the interaction. However, it is not known if these differential effects are dependent on 5-HT receptor specificity. This relationship between 5-HT receptor subtype and aggression in *T. dalmanni* was investigated in the current study.

The extent to which 5-HT modulates discrete aggressive behaviors in invertebrates may also be influenced by the actions of neuropeptide systems, as shown for vertebrates. For example, lesioning neurons containing tachykinin (Tk) receptors reduced violent attacks in rats but left milder attacks unaffected (24). Similarly, high-intensity aggressive behavior during intrasexual contests is elicited by activation of Tk neurons in male *Drosophila* (25). Overlap in function is also seen with neuropeptide Y (NPY) and its invertebrate homolog neuropeptide F (NPF), which decrease frequency of high intensity aggression in mice and *Drosophila*, respectively (12, 22). These findings imply evolutionarily conserved roles for these neuropeptides in determining aggression intensity and conflict escalation.

Moreover, NPY has been shown in mammals to exert its effects by modulating 5-HT activity (22), and conversely, 5-HT directly influences NPY release in brain regions that mediate aggression (26–29). However, unlike mammals, 5-HT and NPF pathways appear to act independently in regulating *Drosophila* aggression (10). Receptors for Tk are located on both 5-HT and non-5-HT neurons within the mammalian 5-HT cell body region (dorsal raphe), and can modulate neuronal firing and 5-HT release in terminal regions (30). In contrast, Tk and 5-HT do not appear to be co-localized to the same neurons in the majority of invertebrates (31–37), and while they have been shown to exert similar excitatory effects on crustacean cardiac and gut ganglia (38,39), it is unclear whether Tk and 5-HT influence each other to regulate invertebrate aggression. Combined, these findings highlight the need to consider multiple discrete measures of aggressive behavior (e.g., frequency and intensity plus contest duration) along with activity of other neuromodulators when determining the role of 5-HT as a whole.

Finally, a seemingly more troubling limitation in our knowledge of how 5-HT mediates aggressive behavior is the lack of studies conducted with female invertebrates. Although several studies have described neural pathways involved in aggression and related behaviors, the almost exclusive use of male subjects precludes determining whether these are sex-specific (for review see 40). Given the stark contrast in aggression and related behaviors between the sexes of many invertebrate species, as well as morphological dissimilarities (1), it is reasonable to suggest significant sex differences in underlying neural mechanisms. For instance, in the stalk-eyed fly, *T. dalmanni*, both sexes possess elongated eyestalks and engage in intrasexual aggressive contests over food resources, but unlike for males, the relatively shorter eyestalks of females do not appear to be used for assessment of opponents (41,42). Further, qualitative comparison across studies using either sex suggests females rarely engage in high intensity aggression (e.g., 1,41,42). Pharmacologically increasing 5-HT in a smaller male will override the inhibition of aggression evoked by perception of a larger opponent with longer eyestalks (23), and expression of high intensity aggression by males in size-matched or mismatched contests is increased by 5-HT treatment (2). The sex difference in opponent perception and aggression level, combined with the known effects of 5-HT on these behaviors in males, makes *T. dalmanni* an ideal model species for examining whether insect aggression is modulated by 5-HT in a sex-dependent manner.

Here, we sought to address some of the gaps in knowledge about 5-HT influences on invertebrate aggression by employing a range of complementary experimental approaches using both male and female *T. dalmanni*. Specifically, we determined 1) whether 5-HT receptor subtypes mediate expression of aggressive intensity and contest progression, 2) if there is a direct functional relationship between 5-HT subtypes and the neuropeptides Tk and NPF in regulating aggression, and 3) how these findings differ between sexes. Using a combination of socio-environmental manipulation, RNA interference (RNAi), and pharmacological treatment, we describe sex-specific roles for 5-HT_2_ receptors, Tk and NPF in behavioral expression and how this relates to contest progression and outcome. We also demonstrate a direct effect of 5-HT on neuropeptide expression that is posited to modulate male aggression. Our results reveal the impressive level of conservation with respect to neurochemical mechanisms among species as diverse as vertebrates and invertebrates, and highlights the need to consider multiple factors when determining potential taxonomic differences in how 5-HT mediates aggression.

## 2. Materials and Methods

### Subjects

The sexually dimorphic stalk-eyed fly, *Teleopsis dalmanni*, is native to South East Asia. All flies used in this study were descendants of pupae collected from wild populations in 2012, Gombak Field Station, housed at the University of Maryland, College Park. Individuals were housed communally in cages (45 cm x 22 cm x 19 cm) on a 12-h light:dark cycle with free access to food, water, and mates. Sexually mature flies in this study were briefly anesthetized using ice and eye span measured to the nearest 0.01 mm using cellSens standard software (Olympus). Size-matched individuals were determined as being < 1% difference in eye span. The high correlation of eye span to body size in this species makes eye span an accurate representation of body size (43, 44). Predetermined opponents were given identifying paint marks between their thoracic spines and housed in separate communal enclosures prior to surgeries and fight contests. Isolated flies were housed individually for 7 days prior to fight contests.

### Drug administration

Flies selected for 5-HT manipulation were fed sterilized, pureed sweet corn containing either 3 g of 5-hydroxy-_L_-tryptophan (5-HTP; H9772; Sigma, St. Louis, MO) in 100 mL media or vehicle/100 mL media for 4 days as previously described (14, 15). Specifically, the vehicle media contained 100 mL of corn, 1 mL of methylparaben (Ward’s Science, Rochester, NY) as a mold inhibitor (Wilkinson 1993), and 25 mg of ascorbic acid (Sigma, St. Louis, MO) to act as a stabilizer (12).

### Forced-fight paradigm and behavioral analysis

Opponent matchups for the experiments described below are as follows: socially isolated vs socially reared, 5-HT_2_ siRNA vs vehicle siRNA, and 5-HTP treated vs vehicle treated. Using previously published methods (e.g., 15), size-and sex-matched opponents were placed in an arena (11 cm x 6.5 cm x 5 cm) containing a glass wall and ceiling, for filming, and a removable cardboard barrier that separated the individuals. Flies were starved for 12 hours prior to the contest to increase the incentive to fight over a piece of pureed corn that was placed in the arena center immediately following barrier removal. All contests that occurred during 10 mins of fighting were scored using the behavioral scoring software, JWatcher (UCLA). Scored behaviors for each individual were based on an existing ethogram adapted from Egge et al. (45). Specific behaviors fall into three categories: contest initiation, escalation, and termination. Contest initiations were determined by one opponent approaching the other, which ultimately results in an aggressive behavioral exchange. Only one opponent was awarded an initiation per contest, none were awarded if there was ambiguity in determining the initiator. Escalations of fights were determined by high-intensity (HI) physical contact behaviors (see 15). Termination and subsequent winners of contest were determined by retreats, with the individual with the fewest number of retreats after the 10 minute fighting period deemed the winner.

### RNA isolation and quantification

RNA isolation was conducted from dissected whole-brain tissue using the Direct-zol™ RNA MiniPrep (ZYMO Research, Irvine, CA) according to the manufacturer’s instructions. Isolated RNA was reverse transcribed using the Maxima First Strand cDNA synthesis kit (Thermo Fisher Scientific, Waltham, MA). Quantitative PCR was performed by an Applied Biosystem’s StepOne machine (Foster City, CA) using TaqMan master mix (Applied Biosystems), the primer-probe pair for the endogenous control, Gapdh, and one of the following target gene primer-probe pairs: 5-HT_1A_, 5-HT_2_, 5-HT_7_, SERT, Tk, or NPFr (Integrated DNA Technologies, Coralville, Iowa; Table 1). We intentionally focused on the Tk ligand itself and not the receptor due to ambiguity in the specific receptor responsible for baseline aggression in dipteran flies (24). Relative quantification (RQ) of the PCR product was conducted using the comparative C_T_ method (46). Values were presented as target gene expression relative to the endogenous control Gapdh).

**Table 1:**
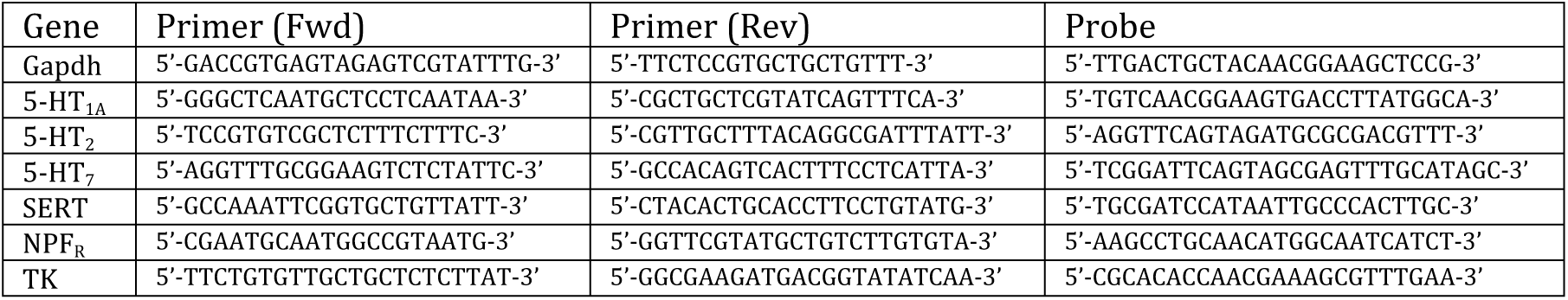
The primer-probe pair for the endogenous control, Gapdh, and the targeted genes.

### RNAi design and administration

Results from the RNA quantification revealed that expression of 5-HT_2_ receptors in male flies was approximately half that seen in females, while 5-HT_1A_ expression was 1.5 times higher in males (see Results and Fig. 1). More aggressive isolated male flies also had lower 5-HT_2_ expression than their socially-housed counterparts, while no difference was seen in 5-HT_1A_ receptors (Fig. 3). Studies in male *Drosophila* have indicated that 5-HT_2_ receptors have a specific inhibitory effect on high intensity aggression, while 5-HT_1A_ receptors promote low intensity (non-contact) behaviors (19). In addition, sex differences in *T. dalmanni* intrasexual aggression predominantly relate to lack of high intensity behaviors in females, which have lower 5-HT_2_ receptor expression than males (Fig. 1). Therefore, we conducted loss-of-function studies targeting the 5-HT_2_ rather than the 5-HT_1A_ receptor for examining how 5-HT receptor subtype modulates aggression and neuropeptide activity in this species.

**Fig. 1.**
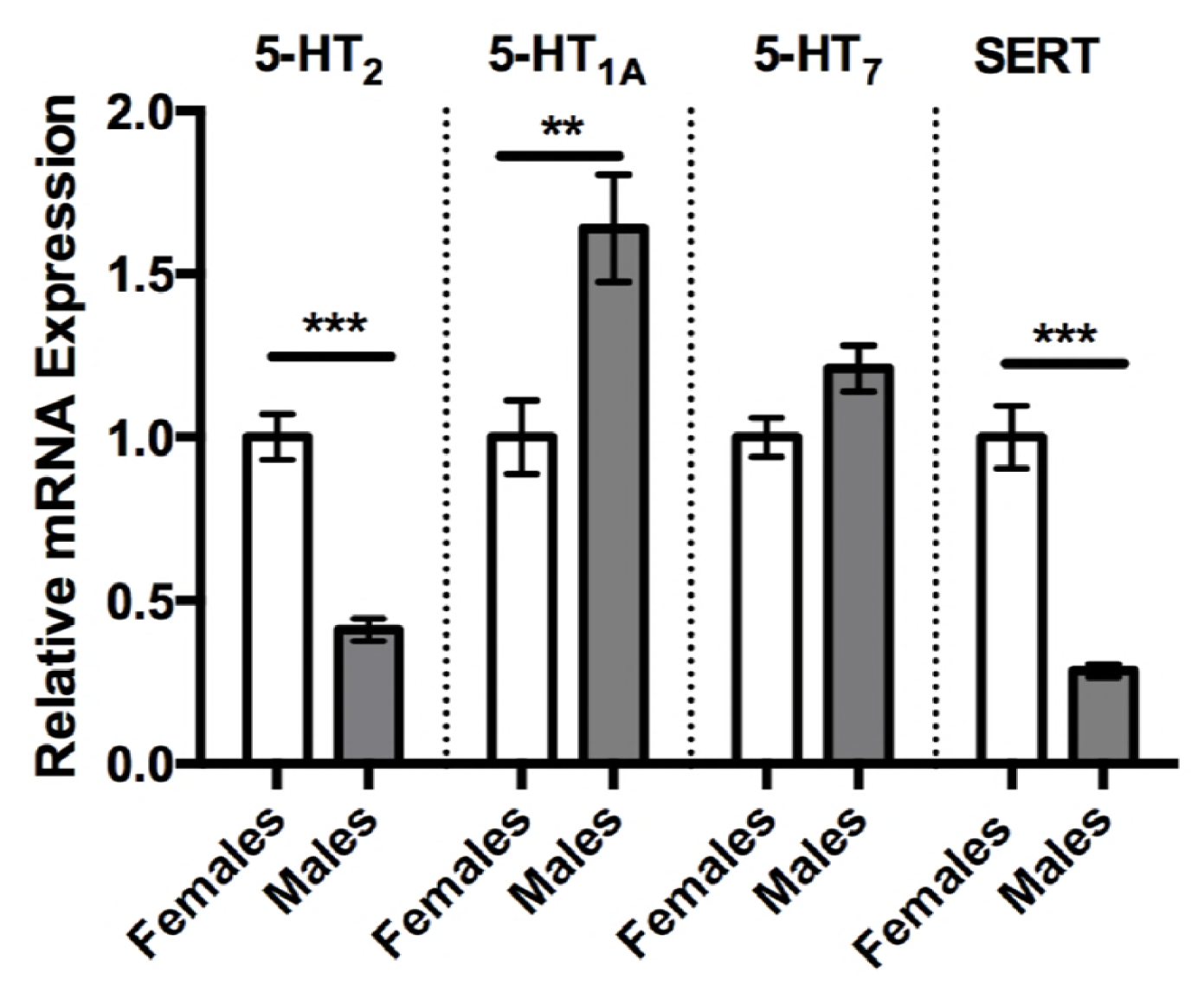
Relative mRNA expression between normally reared adult male and female stalk-eyed flies. Males have a significantly lower expression of 5-HT_2_ (0.41 ± 0.03, n = 9 vs 1.0 ± 0.002, n = 9) and SERT (0.284 ± 0.02, n = 9 vs 1.0 ± 0.001, n = 9) but significantly higher expression levels of 5-HT_1A_ (1.64 ± 0.17, n = 12 vs 1.0 ± 0.001, n = 9) compared to their female counterparts. There was no statistical difference in expression levels of 5-HT_7_ between males and females. Values are normalized to female expression levels and presented as mean ± SEM. (Two-tailed, Student’s *t*-test; *p* < 0.05^*^, *p* < 0.01^**^, *p* < 0.001^***^).

Dicer-substrate siRNAs (short interfering RNA) used in this study were synthesized by IDT (Integrated DNA Technologies, Coralville, IA). Probes targeting 5-HT_2_ were generated using cDNA sequences of *T. dalmanni*. Probes targeting green fluorescent protein (GFP) were generated from cDNA sequences of *Aequorea coerulescens*. The siRNA was coated (1:1 ratio) in the transfection reagent polyethylenimine (PEI) to aid in membrane penetration and stabilization. Injections were administered in the head cavity through the base of the proboscis. The brain was not penetrated with the needle.

Males and females were randomly selected from a social cage and size-and sex-matched to an opponent. Treatment groups were allocated randomly, with experimental males and females being injected with 100 nL of 5-HT_2_ siRNA solution and control flies injected with 100 nL of GFP siRNA solution. All subjects were returned to their previous housing conditions and allowed to recover for 48 hours with free access to food and water. Flies were then placed in the forced-fight paradigm described above.

### 5-HT quantification

Quantification of brain 5-HT was conducted using high-performance liquid chromatography (HPLC) with electrochemical detection as previously described (14). Briefly, whole brains were dissected and frozen in 60 µL of acetate buffer containing the internal standard α-methyl-dopamine (Merck) and stored at −80°C. Samples were thawed and centrifuged at 17,000 rpms. A portion of the sample (45µL) was injected into the chromatographic system and the amines were separated with a C_18_ 4 µm NOVA-PAK radial compression column (Waters Associates, Inc. Milford, MA) and detected using an LC 4 potentiostat and a glassy carbon electrode (Bioanalytical systems, West Lafayette, IN). The CSW32 data program (DataApex Ltd., Czech Republic) calculated monoamine concentrations based on peak height values that were obtained from standards (Sigma-Aldridge, St. Louis, MO). The remaining sample was solubilized for protein analysis with 60 µL of 0.4 M NaOH (47). The final amine concentrations are expressed as pg amine/ µg protein following appropriate corrections for injection vs preparation volume.

### Statistical analysis

To account for behavioral data not meeting assumptions of normality and equal variance, separate two-tailed Wilcoxon matched pairs signed-rank tests were used to compare frequency of contest initiations, expression of high intensity aggression and contest outcome between treated and control groups. Separate two-tailed Student’s *t*-tests were applied to test for relative differences in mean mRNA expression for 5-HT_1A_, 5-HT_2_, 5-HT_7_, SERT, Tk, or NPFr between treatment groups, as well as for differences in mean 5-HT concentrations as quantified by HPLC. Alpha level was set at 0.05 throughout, and analyses performed using Prism (GraphPad Software, Inc; La Jolla, CA).

## 3. Results and Discussion

### Expression of 5-HT_1A,_ 5-HT_2_, 5-HT_7_ receptors and the serotonin transporter in male and female brains

Since 5-HT plays a prominent role in modulating invertebrate aggression (1), including increasing aggressive displays in male *T. dalmanni*, we first measured relative sex differences in mRNA expression of 5-HT_1A_, 5-HT_2_, and 5-HT_7_ receptors and the 5-HT transporter (SERT; Fig. 1). Males had markedly lower expression levels of the 5-HT_2_ receptor subtype (Student’s *t*-test, *p* < 0.001) and SERT (Student’s *t*-test, *p* <0.001) but significantly higher levels of 5-HT_1A_ (Student’s *t*-test, *p* < 0.01; Fig. 1) compared to females. Males and females had similar expression levels of the 5-HT_7_ receptor subtype (Fig. 1).

### Expression of the neuropeptide Tk and the NPF receptor in male and female brains

We also investigated the expression of the NPF receptor in male and female flies, since NPY and the invertebrate analog NPF inhibits aggressive behavior in other animal species (12, 22). We also measured the baseline expression of the neuropeptide Tk, which has been reported to increase aggression in vertebrates and *Drosophila* (24, 25). Baseline relative mRNA expression levels of the NPF receptor (1 ± 0.07, n = 12 vs 0.86 ± 0.05, n = 12; Student’s *t*-test, *p* > 0.05) and Tk (1 ± 0.23, n = 7 vs 0.95 ± 0.17, n = 9; Student’s *t*-test, *p* > 0.05) were similar between females and males, respectfully.

### Increased aggression corresponds with reduced 5-HT_2_ and heightened Tk expression in males but not in females

Social isolation increases aggression in both invertebrate and vertebrate species (19, 48-50). We first tested whether social isolation induced aggression in male stalk-eyed flies. Following 7 days of isolation, males had a higher probability of winning the aggressive confrontation compared to their socially reared opponents (Fig. 2D; 85% vs 15%). Isolated males also initiated significantly more confrontations (Fig. 2A; Wilcoxon matched-pairs signed rank test, *p* < 0.01) and performed more high-intensity (physical) behaviors than their socially-reared opponents (Fig. 2C; Wilcoxon matched-pairs signed rank test, *p* < 0.05). When we repeated this experiment with female flies we found no difference in either initiations or escalations of aggression in socially-isolated females when compared to colony-raised females (Fig. 2B). The lack of alterations in aggressive behaviors in female fights did not allow for easily identified winners, thus the probability of winning the aggressive encounter was not computed.

**Fig. 2.**
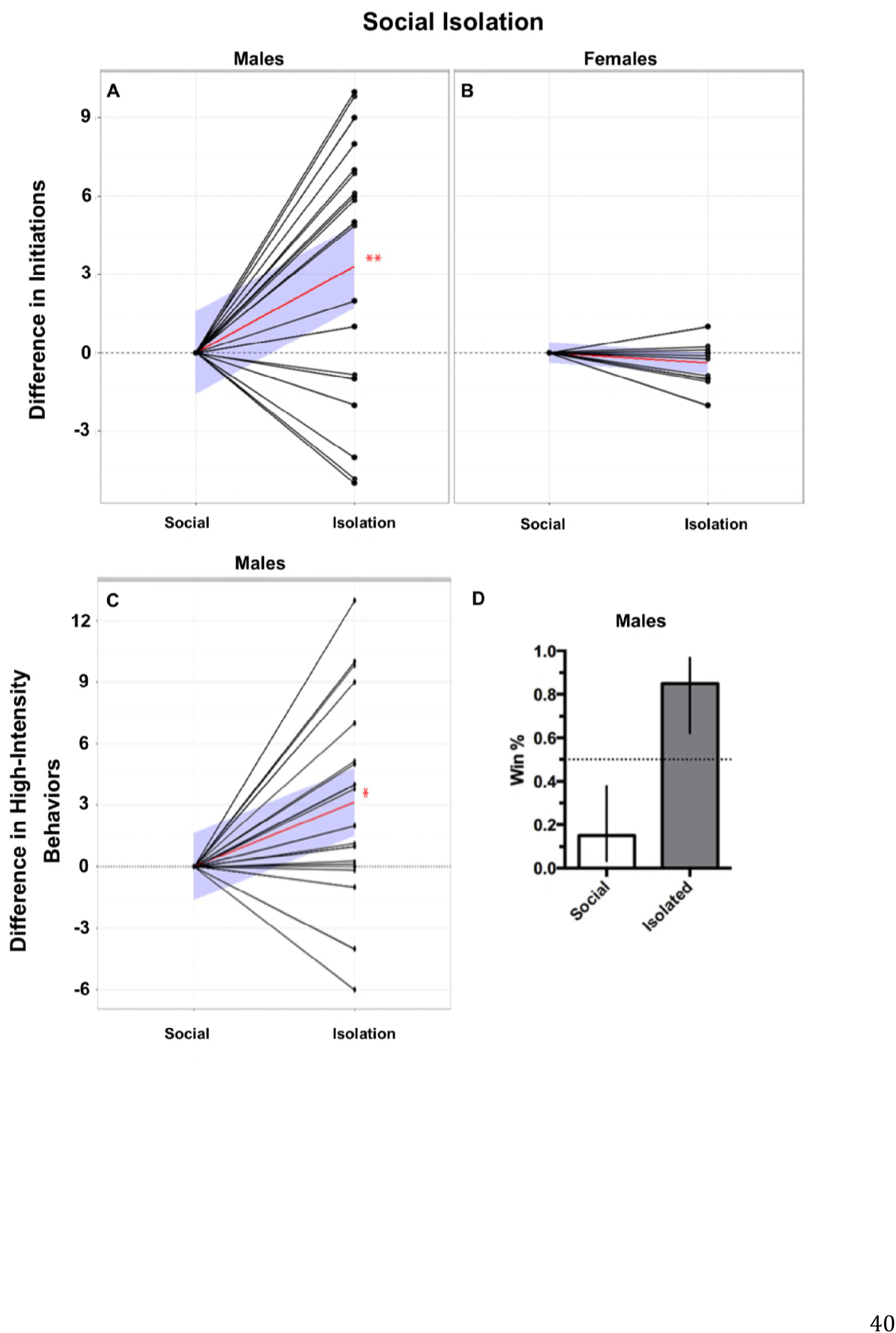
Social isolation increased initiations, escalations, and winning probability in males but not females. A) Males, socially isolated, performed significantly more initiations compared to their socially reared opponents. B) Social isolation had no effect on initiations in female aggressive contests. C) Males, socially isolated, performed significantly more high-intensity aggressive behaviors compared to their socially reared opponents. Behavior values are normalized to control opponents with connecting black lines representing opponents. Red lines represent the mean slope of the differences in behaviors performed between paired opponents and the blue shaded areas represent the 95% confidence interval. (Two-tailed, Wilcoxon matched-pairs signed-rank test; n = 20, *p* < 0.05^*^, *p* < 0.01^**^, *p* < 0.001^***^). D) Males that were socially isolated had a significantly higher winning percentage compared to their socially reared opponents (85% vs 15%; error bars represent 95% confidence interval).

**Fig. 3.**
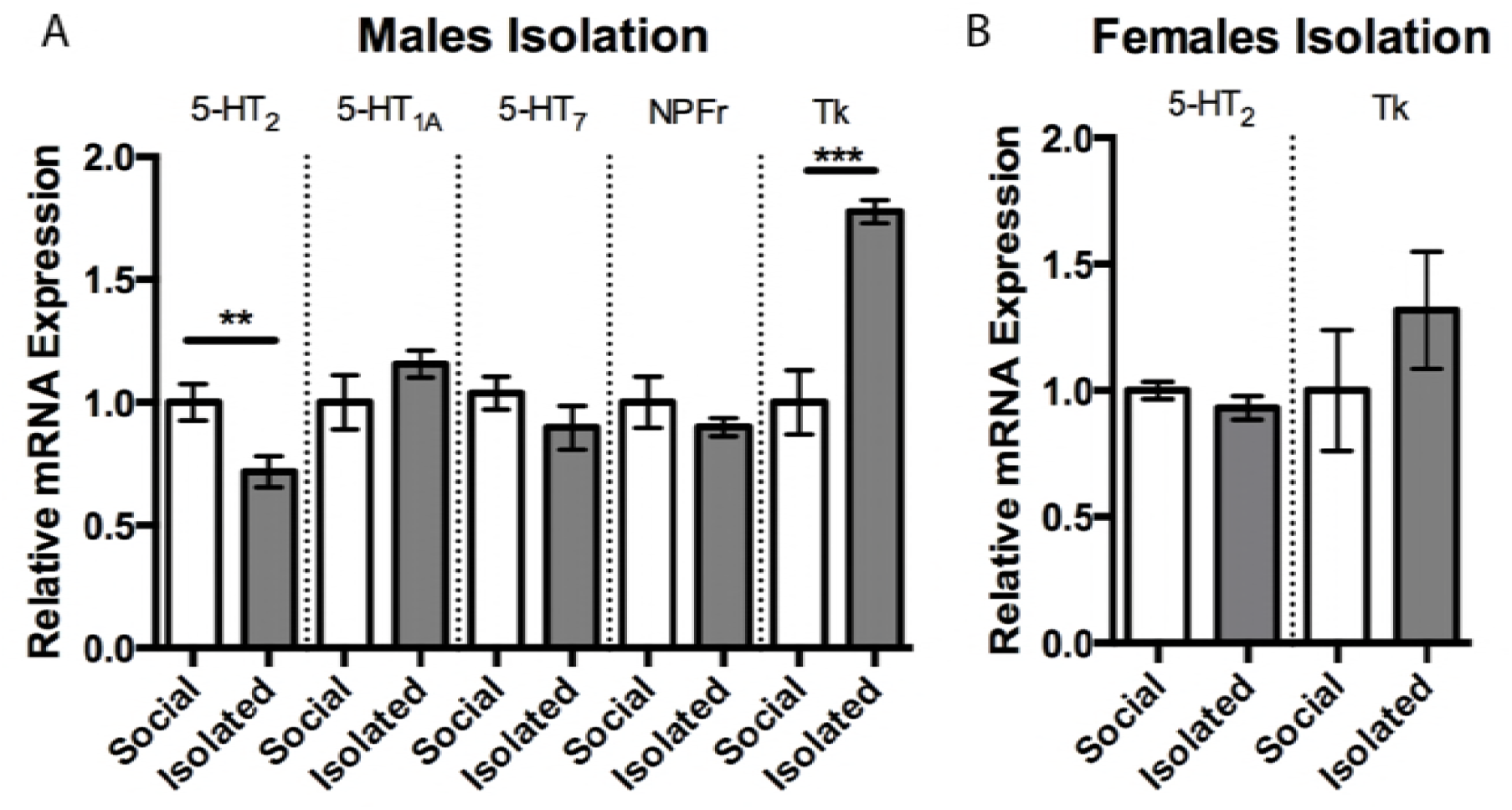
Social isolation reduced 5-HT_2_ expression and increased Tk expression in males but not females. A) Expression of 5-HT_2_ reduced ˜28% in males following social isolation (1.0 ± 0.074, n = 9 vs .72 ± 0.06, n = 12) whereas Tk expression was increased ˜77% compared to socially reared males (1.0 ± 0.13, n = 9 vs 1.77 ± 0.04, n = 6). B) No change in expression value of 5-HT_2_ or Tk was measured in females following social isolation. Similar to males, females did not show differences in 5-HT_1A_, 5-HT_7_, or NPFr expression levels (data not shown). Values are normalized to socially reared expression levels and presented as mean ± SEM. (Two-tailed, Student’s *t*-test; *p* < 0.05^*^, *p* < 0.01^**^, *p* < 0.001^***^).

Next, we measured brain 5-HT_2_ and Tk expression to determine if expression levels covaried with the display of increased aggression. In male flies raised in social isolation, there was a decrease in 5-HT_2_ expression relative to colony-raised flies (Fig. 3; Student’s *t*-test, *p* < 0.01). In contrast, Tk expression was markedly increased in brains obtained from socially isolated male flies relative to colony-raised males (Fig. 3; Student’s *t*-test, *p* < 0.001). Expression of 5-HT_1A_, 5-HT_7_, and NPFr were not different between groups (Fig. 3). In contrast, when the experiment was repeated in females, we found expression levels of 5-HT_2_ and Tk in socially isolated flies did not differ significantly from values obtained in colony-raised flies (Fig. 3). Similar to males, females did not show differences in the expression of brain 5-HT_1A_, 5-HT_7_, or NPFr following isolation (data not shown).

### 5-HT_2_ knock-down increases aggression in males but not in females

To functionally test the role of 5-HT_2_ in aggression, we designed siRNA to selectively knock-down the receptor subtype. Intracranial injections of 5-HT_2_ siRNA reduced 5-HT_2_ receptor expression by approximately 30% 48 hours after the injection compared to control-injected individuals (siRNA generated to target GFP; Fig. 4; Student’s *t*-test, *p* < 0.05). Importantly, the designed 5-HT_2_ siRNA was specific, as it did not affect the expression of the closely related 5-HT_1A_ receptor (Fig. 4).

**Fig. 4.**
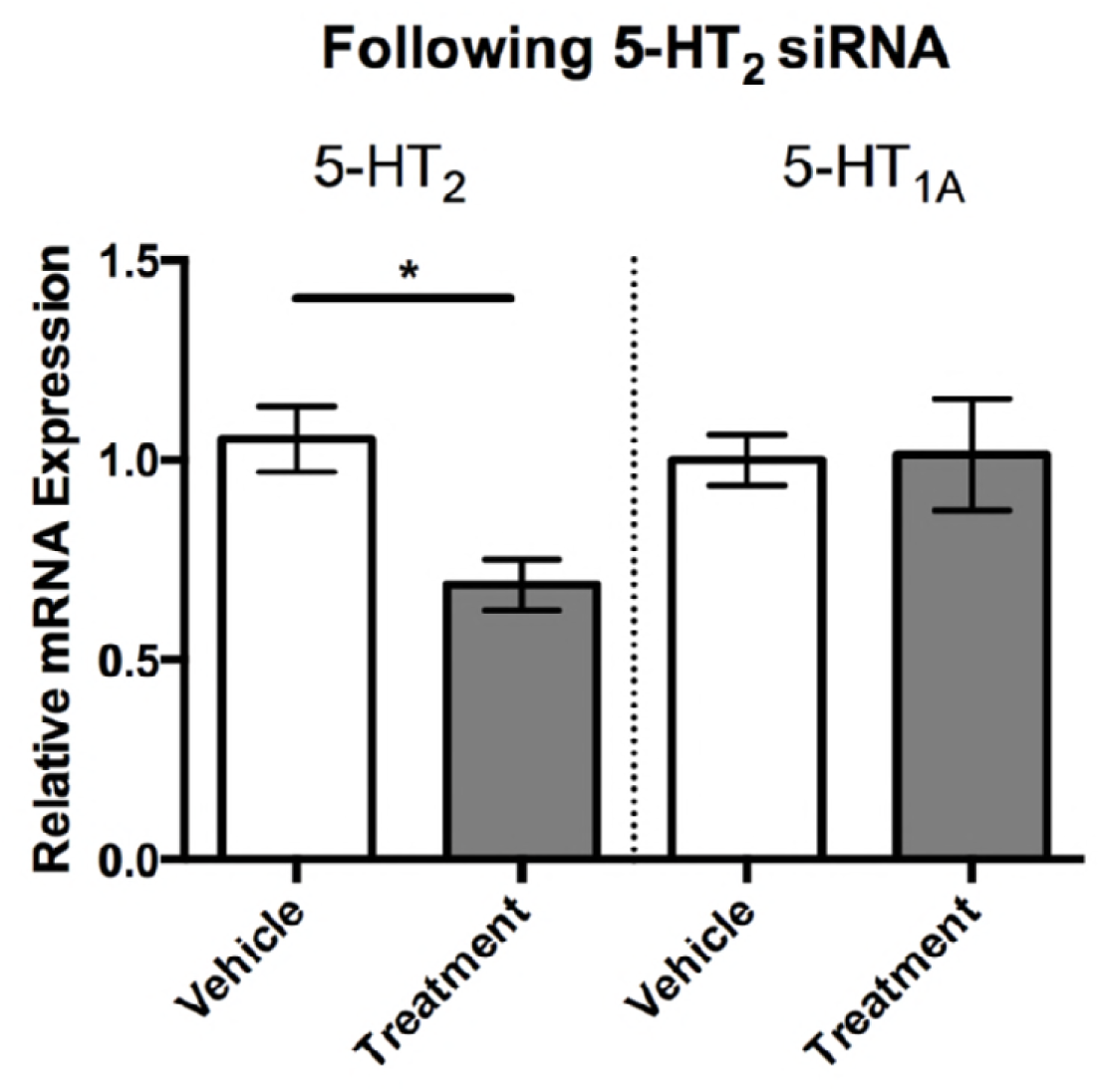
Individual flies injected with 5-HT_2_ siRNA had significantly reduced 5-HT_2_ expression levels compared to vehicle injected siRNA (1.0 ± .09, n = 9 vs 0.68 ± 0.06, n = 6). 5-HT_2_ siRNA did not affect expression levels of the closely related 5-HT_1A_ receptor. Values are normalized to vehicle injected individuals and presented as mean ± SEM. (Two-tailed, Student’s *t*-test; *p* < 0.05^*^, *p* < 0.01^**^, *p* < 0.001^***^).

Socially-reared males injected with 5-HT_2_ siRNA initiated more confrontations compared to their vehicle-injected opponents (Fig. 5; Wilcoxon matched-pairs signed rank test, *p* < 0.001). However, high-intensity behaviors in males did not differ between 5-HT_2_ siRNA injected and vehicle-injected opponents (Fig. 5). Female opponents did not differ in either initiations or high-intensity behaviors, despite 5-HT_2_ siRNA-injected females having significantly lower 5-HT_2_ expression levels compared to their vehicle-treated opponents (Fig. 5).

**Fig. 5.**
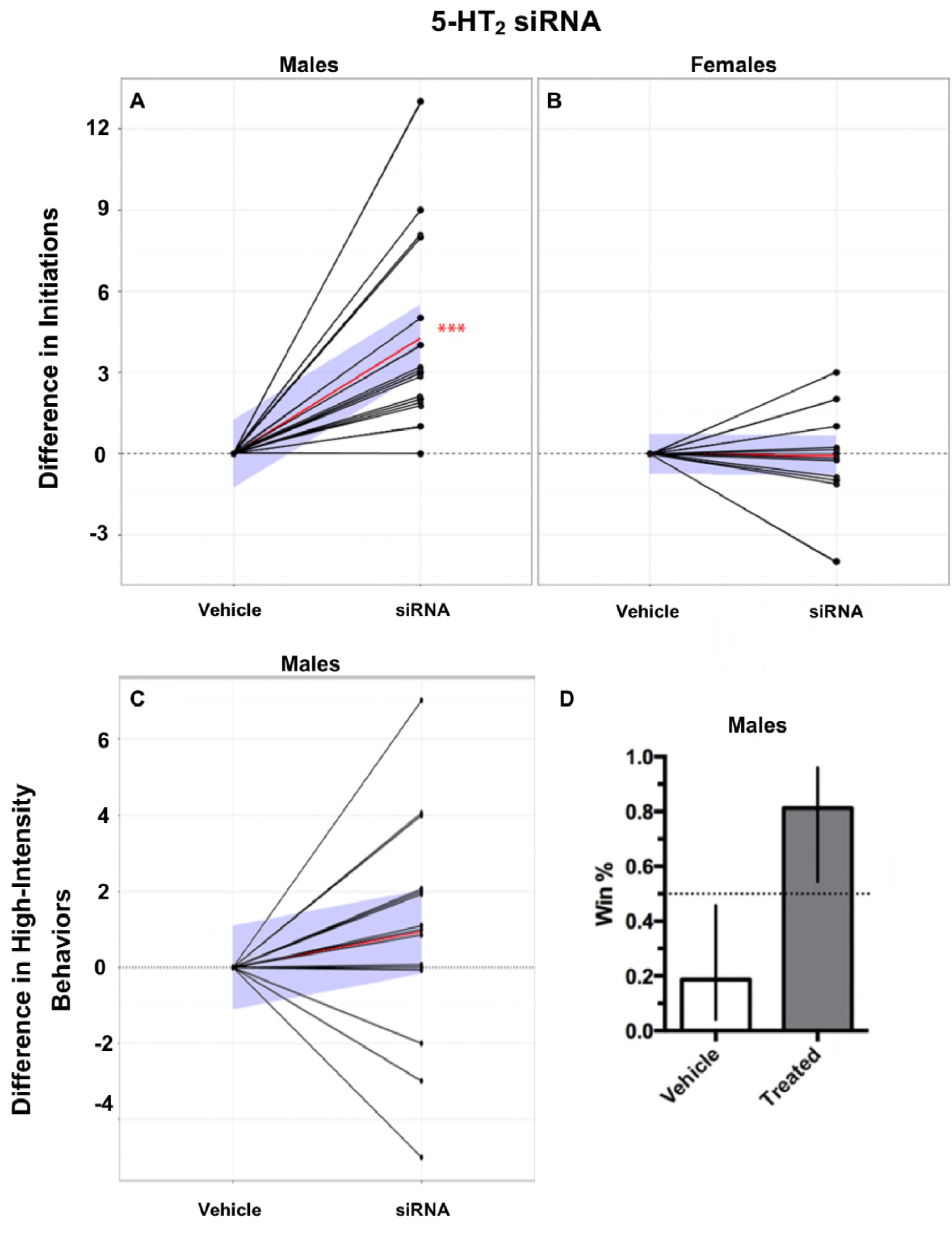
5-HT_2_ siRNA injections increased initiations and winning probability in males but not females. A) Males, injected with 5-HT_2_ siRNA, performed significantly more initiations compared to their vehicle injected opponents. B) 5-HT_2_ siRNA had no effect on initiations in female aggressive contests. C) Males, injected with 5-HT_2_ siRNA, had no differences in high-intensity aggressive behaviors compared to their vehicle injected opponents. Behavior values are normalized to control opponents with connecting black lines representing opponents. Red lines represent the mean slope of the differences in behaviors performed between paired opponents and the blue shaded areas represent the 95% confidence interval. (Two-tailed, Wilcoxon matched-pairs signed-rank test; n = 16, *p* < 0.05^*^, *p* < 0.01^**^, *p* < 0.001^***^). D) Males, injected with 5-HT_2_ siRNA, had a significantly higher winning percentage compared to their vehicle injected opponents (81% vs 19%; error bars represent 95% confidence interval).

### 5-HT_2_ activity modulates NPFr expression in males but not females

To probe for potential interactions between 5-HT and neuropeptides in modulating aggression in males, we next measured brain NPFr and Tk expression levels following the siRNA-induced reduction in 5-HT_2_. Reducing expression of 5-HT_2_ using siRNA significantly reduced NPFr expression in males (Fig. 6; Student’s *t*-test, *p* < 0.01). When this experiment was repeated in females, our results showed that reduction of 5HT_2_ receptors did not significantly affect NPFr expression (Fig. 6). Expression of Tk did not differ in males or females treated with 5-HT_2_ siRNA (Fig. 6).

**Fig. 6.**
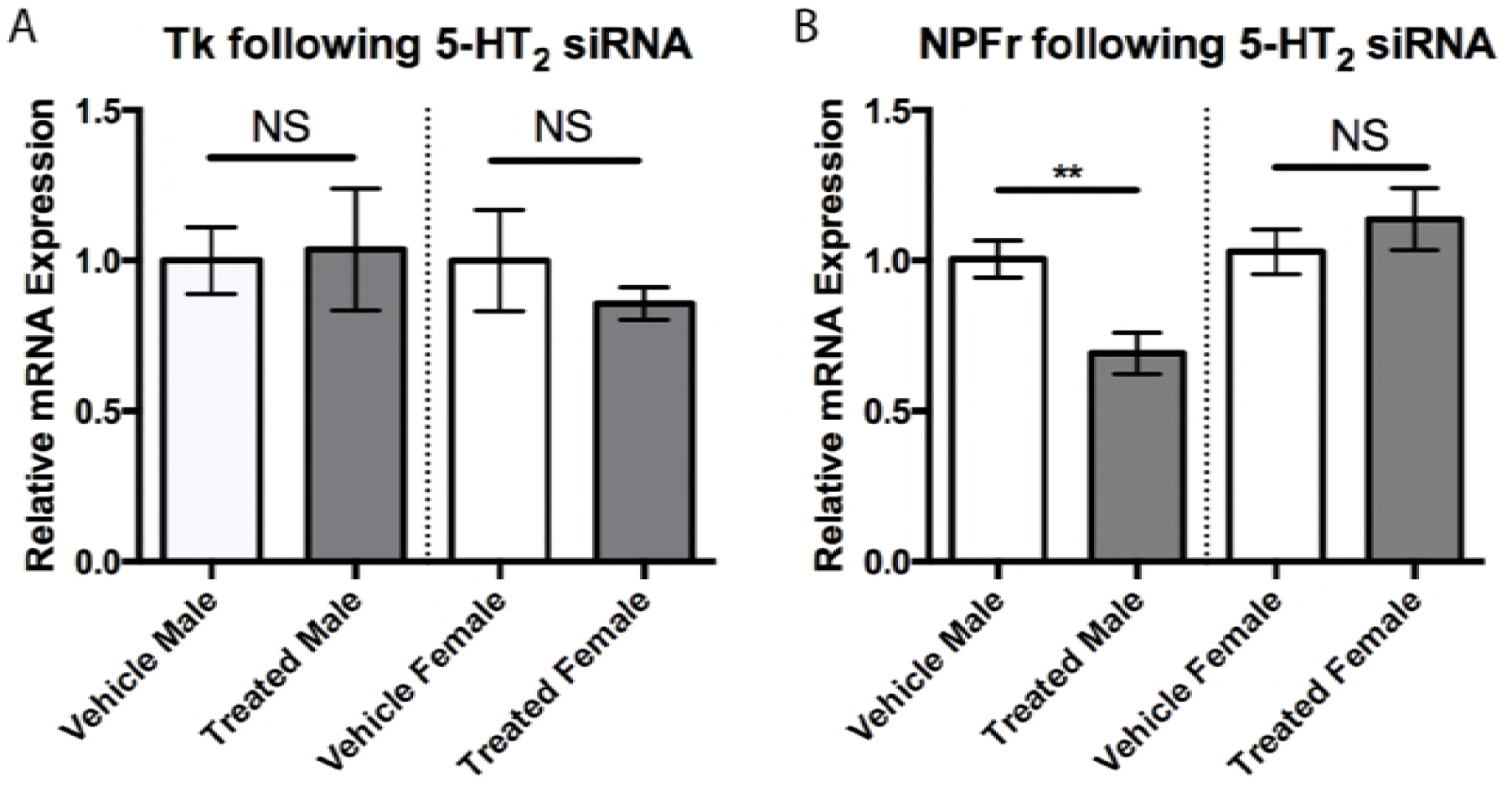
A) Injections of 5-HT_2_ siRNA did not alter expression levels of Tk in either males or females compared to their vehicle injected counterparts. B) Injections with 5-HT_2_ siRNA reduced NPFr expression in males (0.69 ± 0.07, n = 9 vs 1.0 ± 0.06, n = 12) compared to vehicle injected individuals but not in females. Values are normalized to female expression levels and presented as mean ± SEM. (Two-tailed, Student’s *t*-test; *p* < 0.05^*^, *p* < 0.01^**^, *p* < 0.001^***^).

### Elevating 5-HT concentrations increases Tk expression in males but not females

We have previously reported an increase in high-intensity aggressive behaviors in male stalk-eyed flies following administration of the 5-HT metabolic precursor, 5-HTP (15). When we repeated this study in female stalk-eyed flies, we found that 5-HTP-treated individuals do not exhibit a significant difference in aggressive behaviors compared to their untreated opponents, despite having significantly elevated brain 5-HT concentrations (18.44 ± 2.99, n = 18 vs 9.8 ± 0.6, n = 19; pg/μg protein; Student’s *t*-test, *p* < 0.01). In fact, the observation of a fight containing a single high-intensity behavior was so rare that accurate measurements couldn’t be taken. We next explored whether this sex-difference in the expression of high-intensity behaviors following 5-HTP administration might be related to changes in Tk expression. Indeed, males treated with 5-HTP had significantly higher Tk expression compared to vehicle-treated males (Fig. 7; Student’s *t*-test, *p* < 0.001). In contrast, females treated with 5-HTP did not have a significant difference in Tk expression when compared to controls (Fig. 7A). Additionally, expression of NPFr was also significantly raised in males following 5-HTP treatment but not in females (Fig. 7B; Student’s *t*-test, *p* < 0.001). We did not find significant differences in the expression of brain levels of 5-HT_1A_, 5-HT_2_, or 5-HT_7_, respectively, in either males or females treated with 5-HTP when compared to vehicle-treated controls.

**Fig. 7.**
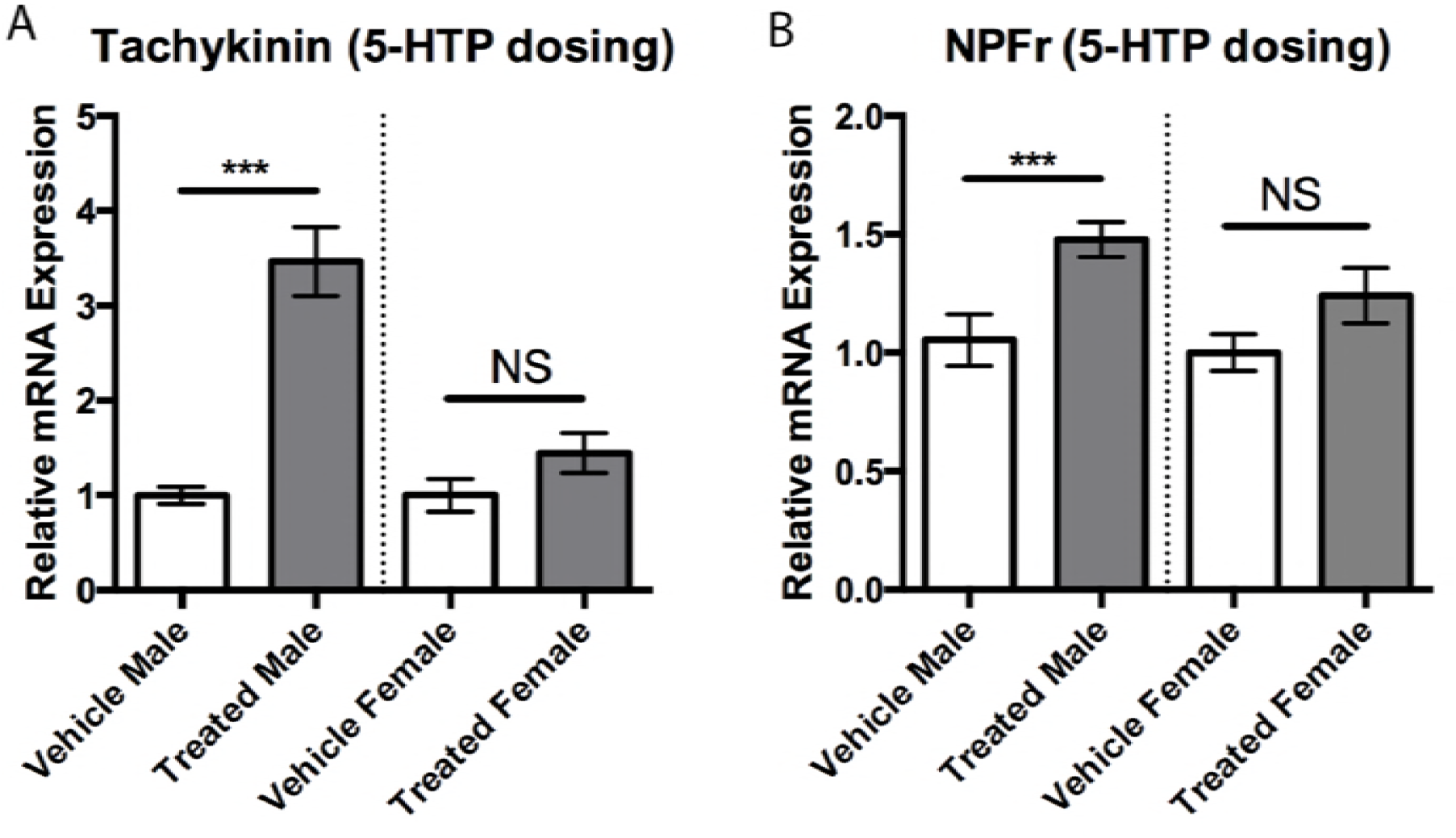
A) Treatment with 5-HTP increased expression levels of Tk in males compared to vehicle treated individuals (3.46 ± 0.36, n = 12 vs 1.0 ± 0.09, n = 9) but not in females. B) Treatment with 5-HTP increased expression levels of NPFr in males compared to vehicle treated individuals (1.48 ± 0.07, n = 12 vs 1.05 ± 0.11, n = 9) but not in females. Values are normalized to female expression levels and presented as mean ± SEM. (Two-tailed, Student’s *t*-test; *p* < 0.05^*^, *p* < 0.01^**^, *p* < 0.001^***^).

In these studies, we used the sexually dimorphic stalk-eyed fly as a model to study how aggression is mediated by both 5-HT and neuropeptides. This model offers several advantages for obtaining a better grasp, both functionally and evolutionarily, of the origin and regulation of this complex behavior across taxa and sexes. First, flies contain fewer neurons and neuronal connections compared to vertebrates while simultaneously maintaining the relevant behaviors. Additionally, the monoaminergic and peptidergic systems involved in these behavioral processes are highly conserved across taxa (18, 24, 25, 51–54). This is not surprising given the fitness benefits and selection pressures of engaging in aggressive conflicts in most living animal species. Finally, the extreme differences in both morphology and behavior between male and female stalk-eyed flies provides a valuable perspective of the proximate mechanisms that have been acted upon by selection forces to result ultimately in sex-specific aggression.

Here, we found that male and female flies raised in social colonies differentially express central 5-HT_1A_-and 5-HT_2_-like receptor subtypes and SERT. Specifically, males show higher expression of 5-HT_1A_ receptors but lower expression of 5-HT_2_ receptors and SERT compared to females. In contrast, 5-HT_7_ receptor expression was equivalent in both sexes. Naturally reduced SERT in males would presumably result in lower 5-HT clearance and corresponding increases in extracellular 5-HT. Combined with the positive relationship between experimentally enhanced 5-HT and male aggression shown by ourselves (14, 15) and others (12, 13, 19), this points to inherently greater serotonergic activity as an explanation for observed sex differences in stalk-eyed fly aggression. Further, our findings suggest this is principally mediated through a balance of signaling in favor of 5-HT_1A_ versus 5-HT_2_ receptor activation.

To test this, we investigated the effects of social isolation on aggression and expression of serotonin receptor subtypes in male and female stalk-eyed flies. Previous studies indicate that social vertebrates and invertebrates that are either reared in isolation or exposed to periods of isolation exhibit increases in aggression to conspecifics (19, 48-50, 55). In line with this, we found that socially isolated male stalk-eyed flies increase conflict initiations, fight escalation, and have a higher probability of winning a fight. This result suggests that the absence of socialization in male stalk-eyed flies may impair detection of social cues that dampen normal aggressive responses and prevent the escalation of a fight to potentially injurious intensities. The finding that isolated males exhibit higher levels of aggression is similar to isolation effects on *Drosophila* behavior (19). In contrast, while isolation also increases aggression in female *Drosophila* (56), socially reared and socially-isolated females in our study exhibited similar levels of initiations and escalations in aggressive behavior. When we evaluated the expression of 5-HT-like receptors, there were no differences in either 5-HT_1A_ or 5-HT_7_ receptors in either sex when compared to socially reared flies. However, the expression of 5-HT_2_ receptors decreased in males but not females when compared to socially reared controls. These results differ from earlier work in *Drosophila* which showed that socially-isolated males had decreased expression of 5-HT_1A_-and increased expression of 5-HT_2_-like receptor mRNA (19). These differences may be species specific, but more likely may be related to the isolation protocol employed. In the *Drosophila* study, the flies were isolated 2 to 3 days post-eclosion and kept in constant light, whereas we isolated sexually mature flies for 7 days and used a 12:12 light dark cycle. Regardless, the isolation-induced reduction in 5-HT_2_ receptors seen here would allow for greater activation of 5-HT_1A_ receptors even in the absence of altered expression of the latter subtype. Together with our behavioral findings, this further supports the notion that decreased and increased activation of 5-HT_2_ and 5-HT_1A_ receptors, respectively, is a likely substrate for heightened aggression following either social isolation or enhanced serotonergic activity.

In addition to measuring 5-HT-like receptor subtypes, we also evaluated the expression of Tk and NPF receptors in socially-isolated and socially reared flies. Neither NPF receptors nor Tk in females was affected by social isolation, and expression of NPF receptors did not differ with rearing condition in males. In contrast, the expression of Tk was markedly increased in socially-isolated males. Recent work in *Drosophila* has shown that neurons expressing Tk regulate male-male aggression (25), suggesting the increase in Tk found in socially-isolated male stalk-eyed flies likely contributes to the increase in aggression displayed by these insects, similar to what is observed in vertebrate species (57). Further, the finding that only males showed both increased Tk and reduced 5-HT_2_ receptors implicates the combination of these factors in determining sex-specific changes in aggression caused by social isolation.

The decreased expression of 5-HT_2_-receptors, increased Tk expression and heightened aggression following social isolation led us to investigate if there was a direct relationship among these factors, which we tested by genetic knockdown of 5-HT_2_ receptors. In line with our prediction, males exhibited increased aggression with 5HT_2_ receptor knockdown, suggesting that serotonergic activation of the 5HT_2_-like receptor normally acts to dampen aggression. This result is consistent with earlier work in *Drosophila,* which showed that treatments with 5-HT_2_ agonists decreased aggression (19), and further implicates the balance between 5-HT_1A_ and 5-HT_2_ signaling in determining expression of aggressive behavior. Interestingly, 5-HT_2_ knockdown had no effect on male Tk expression, suggesting that the increase in Tk observed in socially isolated males is independent of the decrease in the expression of the 5-HT_2_ receptor subtype. While Tk was not affected by 5-HT_2_ knockdown, there was a significant reduction in expression of NPF receptors in male brains. In *Drosophila*, NPF decreases aggression (12), hence the decreased expression of NPF receptors (and subsequent dampening of NPF signaling) seen in our study may contribute to the increased aggression exhibited by male stalk-eyed flies following knockdown of the 5-HT_2_-like receptor. Work using genetically modified male *Drosophila* suggests that 5-HT and NPF function independently to modulate aggression in an additive manner (12). Similarly, increased aggressive behavior displayed by neuropeptide Y1 receptor-knockdown mice is normalized by pharmacological activation of 5-HT_1A_ receptors (22), implying independent actions of 5-HT and NPY in mediating vertebrate aggression. However, other behaviors mediated by central NPY activity in rodents, such as feeding and helplessness, have been shown to be directly linked with 5-HT_2_ receptor function (58, 59). Our results suggest that aggression in stalk-eyed flies may be similarly modulated by 5-HT_2_ receptor-dependent interactions between 5-HT and NPF signaling, in contrast to the dissociated effects of 5-HT and NPF on aggression as shown in mice and *Drosophila*. Combined, it is tempting to speculate that activation of 5-HT_2_ receptors in *T. dalmanni* may directly increase NPF to act as a brake on expression of male aggression.

While only NPF receptor expression was affected by 5-HT_2_ knockdown, male flies showed increases in both Tk and NPF receptor expression following 5-HTP administration. The finding that enhancing 5-HT levels can increase NPF receptors is reminiscent of neurochemical interactions in the mammalian hypothalamus, where HT can directly enhance release of the vertebrate homolog NPY (26, 27), and suggests similar mechanisms in stalk-eyed flies. The means by which 5-HT may be acting to increase Tk in male flies is not clear, given the lack of neuronal co-localization of Tk-like peptides and 5-HT in most invertebrates (31–37). However, 5-HT neurons do appear to synapse on to Tk-immunoreactive terminals in the brain of the desert locust, *Schistocerca gregana* (34), suggesting a direct functional relationship via synaptic contact that may also be present in *T. dalmanni*.

Notably, the effects of 5-HT_2_ receptor knockdown on aggression and NPFr were specific to males, with treated females showing no change either aggressive behavior, Tk or NPFr compared to vehicle-treated females. Similarly, 5-HTP treatment, which reliably increases male aggression (14, 15), had no effect on female aggression, despite causing an increase in brain 5-HT. Effects of social isolation both on aggression and on expression of 5-HT_1A_ and 5-HT_2_ receptors and Tk were also restricted to males. When taken in conjunction with the fact that males have naturally lower 5-HT_2_ but elevated 5-HT_1A_ receptor expression compared with females, along with reduced SERT, this implies that male aggression is much more tightly linked with elevated serotonergic signaling via specific subtypes. Specifically, we posit that reductions in 5-HT_2_ activation in favor of 5-HT_1A_ function may form the mechanistic basis of sex differences in regulation of aggression in this species. Further, the effects of altered 5-HT activity in promoting male aggression are in turn modulated by Tk and NPF, whereas these neuropeptides appear to play no role in female aggression. Identifying these mechanisms is an important step in elucidating factors driving the evolution of sex-specific behaviors in invertebrates.

Comparing the effects of socio-environmental factors against those of genetic and pharmacological manipulation provided a valuable opportunity to elucidate how discrete components of an aggressive interaction are specifically mediated by monoaminergic and peptidergic activity (Table 2). For example, the increase in both aggression and NPF receptor expression following 5-HTP treatment in male flies seems counterintuitive, since NPF is posited to be inhibitory towards aggression. However, parsing out contest initiation, escalation and outcome across experimental procedures reveals that while 5-HTP administration increases expression of high intensity behaviors, it does not affect initiations (14,15). In contrast, increases in initiations were seen after both social isolation and 5-HT_2_ receptor knockdown, and were associated with either no change or a decrease in NPF receptor expression, respectively. Increases in NPF signaling may therefore only be important in decreasing the motivation to engage in the opening stages of the contest (Fig. 8), but have little to no effect in fight escalation, which differs from the direct suppression of high intensity aggression by this peptide seen in mice (22) and *Drosophila* (12). Similarly, both 5-HTP treatment and 5-HT_2_ receptor knockdown enhance the probability of winning, yet each manipulation results in opposite changes to NPF receptor expression (Table 2), suggesting NPF has no role in determining contest outcome. Stage specific functions for 5-HT_2_ receptors and Tk were also indicated when comparing across experimental manipulations (Table 2), with a primary role suggested for 5-HT_2_ receptors in contest initiations, while Tk serves to modulate high intensity aggression (Fig. 8). However, winning appears to be associated with both reduced 5-HT_2_ receptor expression and heightened Tk (Table 2). Together, findings suggest the motivation to engage in a contest is enhanced by decreased 5-HT_2_ receptor activation, which may be potentiated by reduced NPF signaling. As the contest progresses, reduced 5-HT_2_ activity is maintained but no longer influences behavior, with escalations in intensity driven largely by increases in Tk. The combination of increased initiations and high intensity aggression then serves to increase the probability of winning (Fig. 8).

**Table 2.** Effects of experimental manipulations on expression of 5-HT receptors, neuropeptides and aggressive behavior in male stalk-eyed flies, *Teleopsis dalmanni*.

| Manipulation | 5-HT<sub>1A</sub> | 5-HT<sub>2</sub> | NPF | Tk | Contest<br>Initiations | High<br>intensity<br>aggression | Wins |
| --- | --- | --- | --- | --- | --- | --- | --- |
| Social<br>isolation | No<br>change | ↓ |  | ↑ | ↑ | ↑ | ↑ |
| 5-HT<sub>2</sub><br>knockdown | No<br>change | ↓ | ↓ | No<br>change | ↑ | No<br>change | ↑ |
| 5-HTP<br>treatment | No<br>change | No<br>change | ↑ | ↑ | No<br>change* | ↑* | ↑* |

\*From Bubak et al., 2014b

**Fig. 8.**
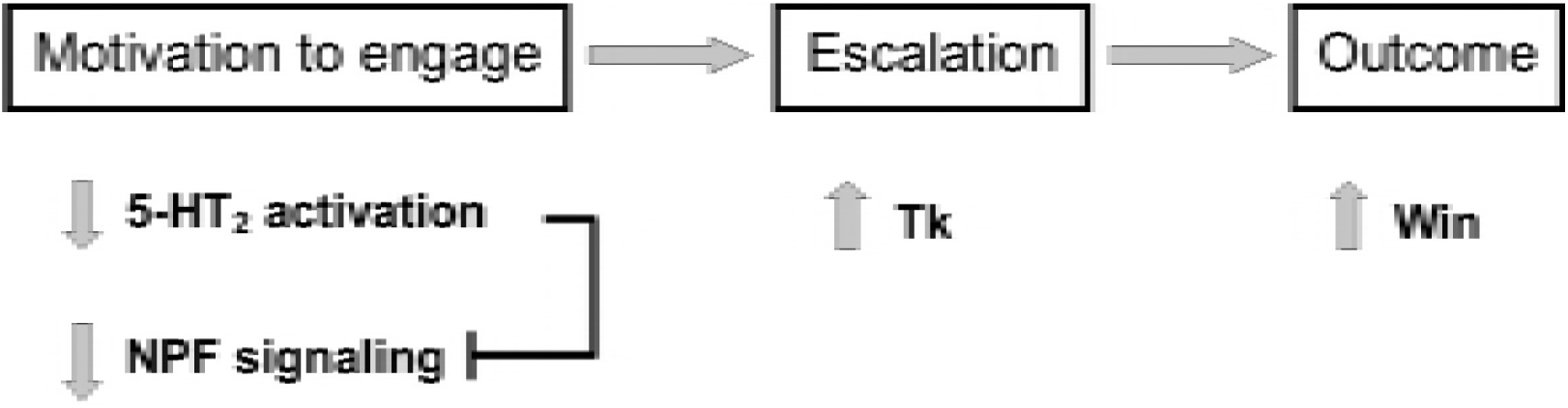
Schematic diagram illustrating proposed relationships between 5-HT / neuropeptide signaling and stages of aggressive conflict in male stalk-eyed flies. The motivation to engage in aggression appears to be promoted by a reduction in 5-HT_2_ receptor activity, which may directly inhibit NPF signaling. Escalation in aggression as the contest proceeds is driven by an increase in Tk, with 5-HT_2_ receptors and NPF no longer influencing behavior. Greater willingness to engage and subsequent expression of high intensity aggression then facilitate the probability of winning.

It is generally presented that 5-HT has a largely inhibitory role in vertebrate aggression, including humans (60–65) while facilitating invertebrate aggression (1, 6, 12–15, 66–68). Here, we used a novel invertebrate model to show a more nuanced role for 5-HT and its receptor subtypes in modulating aggression according to the stage of conflict. Similarly differentiated functions of 5-HT receptor subtypes on the type of aggressive behavior are seen in mice and *Drosophila*, suggesting evolutionary conservation in neurochemical modulation of social behavior. However, it should be noted that our results with stalk-eyed flies point to a primary effect of NPF on contest initiation and not escalation, whereas the reverse pattern is seen in *Drosophila* (12). Our findings also indicate the possibility for a direct functional relationship between 5-HT receptors and NPF signaling in regulating aggression, which does not appear to be present in *Drosophila* (12). Such differences between two Dipteran fly species within the same taxonomic section (Diptera: Schizophora) cautions against generalizing findings from one species across others, but at the same time offers a valuable means of examining mechanisms leading to evolutionary divergence in behavior. We also show that the impact of 5-HT’s interactive role with neuropeptides on discrete stages of aggressive conflict can be inhibitory, stimulatory, or even absent depending on the particular neuropeptide and on the sex of the individual. Thus, stating opposing roles for 5-HT between vertebrates and invertebrates overlooks the intricate and conserved interactive role this monoaminergic system has with other neurochemical systems known to influence aggressive behavior, while also not accounting for effects of sex. Investigating whether these interactions exist and function similarly in a range of animal species undergoing diverse selection pressures will aid in uncovering the origins, and by extension mechanisms, of this complex behavior.

## Acknowledgments

This work was funded by NSF Grants IOS 1256898 (to J.G.S) and IOS 1257679 (to M.J.W.). We thank Dr. Michael Greene for his helpful comments that improved the manuscript.

## Author Contributions

A.N. Bubak wrote the paper and designed and conducted the behavioral experiments. J.G. Swallow contributed to the design of the behavioral experiments and contributed to writing the manuscript. J.D.W. Yaeger and K.J. Renner conducted the monoamine analysis and contributed to writing the manuscript.

